# Software solutions for reproducible RNA-seq workflows

**DOI:** 10.1101/099028

**Authors:** Trevor Meiss, Ling-Hong Hung, Yuguang Xiong, Eric Sobie, Ka Yee Yeung

## Abstract

Computational workflows typically consist of many tools that are usually distributed as compiled binaries or source code. Each of these software tools typically depends on other installed software, and performance could potentially vary due to versions, updates, and operating systems. We show here that the analysis of mRNA-seq data can depend on the computing environment, and we demonstrate that software containers represent practical solutions that ensure the reproducibility of RNAseq data analyses.

---

Reproducibility of results is essential to scientific research. With the generation of diverse and complex biomedical “big data”, the development of computational methods and analysis workflows has become integral to biomedical research. Computational tools are usually distributed as compiled binaries or source code. However, running these binaries or compiling the provided source code requires additional software and depends on the computer environment on the user’s host system. Most importantly, modern biomedical workflows and pipelines consist of multiple applications and libraries, each with their own set of software dependencies. Simply depositing the code in a repository is often <u>not</u> enough to allow a researcher to replicate the analyses. Hence, suites such as Bioconductor^1^, BioPython^2^, Galaxy^3^ and BioPerl,^4^ where the user is assured that the dependencies for the components are properly installed, have become increasingly popular. Although these suites serve an important role, *compatibility* often becomes dependent on the version of the suite that is used. For example, Bioconductor^1^ packages are typically run in the R programming environment, and hence, performance depends on the version of R and other packages installed on the host system. Therefore, published analytical results may not be reproduced if the software is deployed on a computer with different operating systems, alternative configurations, or different versions of installed software dependencies.

The ultimate goal in ensuring reproducibility of bioinformatics analyses is to develop computer software such that an application will give 100% identical results regardless of the host environment. One simple strategy would be to distribute the entire computing environment in the form of a software computer that runs on top of the host system, a configuration called a virtual machine (VM). A VM is a self-contained computing environment that behaves as if it were a separate computer. Therefore, VMs make it possible to export and reproduce the exact environment regardless of the underlying hardware and operating system. While VMs can exactly replicate a given workflow and its dependencies, a separate VM would be needed for each workflow to be shared. This is a poor fit for the modular bioinformatics workflows where separate components are mixed and matched into analyses pipelines. Recently, a popular open-source project, Docker (https://www.docker.com/), has become the *de facto* standard for container software. Docker provides a series of tools to construct, update, distribute and deploy software containers on Windows, Linux, Macintosh and cloud platforms. Docker also has a built-in version control system. It has been shown that Docker containers have only a minor impact on the performance of common genomic pipelines^5^. Our group has proposed solutions to add graphical user interfaces to the command-line based Docker to enhance the usability of containers among the biomedical community^6, 7^.

Here we have addressed whether RNA-seq analyses can be readily reproduced across different computing platforms. RNA sequencing has become the established method to measure genome-wide gene expression^8, 9^. A key step in the analyses of RNA-seq data is the read mapping and expression quantification step in which RNA-seq reads are mapped and aligned to the reference genome, thereby generating a table of read counts across the genome^10^. A common subsequent step is to identify genes that are differentially expressed between different experimental conditions using these count-based transcript expression levels.

We illustrate the challenge of reproducing results in the differential expression inference step. The LINCS (Library of Integrated Network-based Cellular Signatures) Consortium, funded by the National Institutes of Health Common Fund, aims to comprehensively quantify how a variety of cell types respond to pharmacological, environmental, and genetic perturbations^11^. The Drug Toxicity Signature Generation (DToxS) Center, at Icahn School of Medicine at Mount Sinai, is one of six data and signature generation centers within the LINCS Consortium. The goal of the DToxS Center is to provide insight into drug-induced adverse events by generating signatures of how drugs influence mRNA and protein levels, then using these data to build predictive statistical and mechanistic mathematical models. As an initial effort in this direction, DToxS has investigated kinase inhibitors used to treat cancers that may cause cardiotoxicity as an adverse event. In these experiments, drug-induced changes in gene expression in cardiac-derived cells are quantified by mRNA-seq, using a method that employs 3’ end-directed sequencing and tags each mRNA molecule with a Unique Molecular Identifier (UMI) Data are then analyzed using a workflow developed in house.

To determine whether the analysis could be readily reproduced, two standard operating procedures (SOP), describing the generation of gene read counts and the identification of differentially expressed genes,^12^ were implemented by an independent group (see Figure 1 for a summary of the SOP). The analysis workflow includes: (1) mapping sequence fragments onto genes using the Burrows-Wheeler Aligner (BWA)^13, 14^, (2) performing quality control to remove outlier samples and genes using hierarchical clustering analysis, (3) data normalization using the Trimmed Mean of M-values (TMM)^15^, and (4) identification of differentially expressed genes using the edgeR package^16^.As with most bioinformatics workflows, this RNA-seq analysis pipeline consists of multiple components, and each component written in a different language. For instance, the Burrows-Wheeler Aligner (BWA) is a command line tool written in C, whereas edgeR is written in R. Each component that has been written by a different group and in a different language could potentially have different dependencies on other software and settings on the host system. For example, the edgeR Bioconductor package is dependent on both another Bioconductor package, LIMMA,^17^ and on the version of R installed on the host computer.

**Figure 1.**
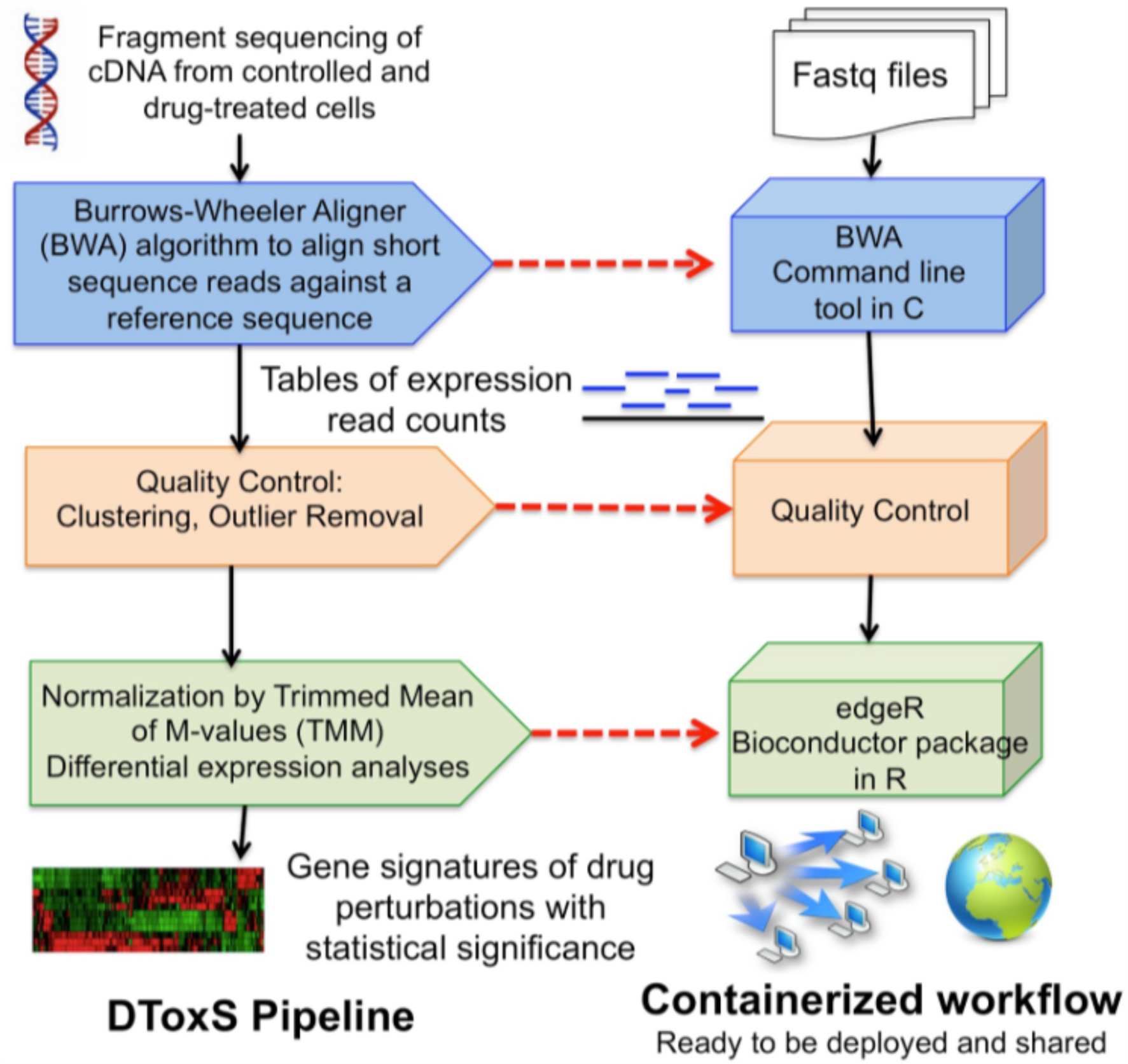
Summary of the RNA-seq SOP from DToX. There are two main steps in the SOP: the sequence alignment step and the differential expression analyses step. A Docker container is created to correspond to each of these steps.

We executed the code and scripts implementing the aforementioned DToxS workflow across multiple operating systems to test the ease of installation and reproducibility of the analysis. The outputs across multiple operating systems and versions of dependencies were compared to find differences that most researchers developing bioinformatics tools might not be aware of. The results from running the DToxS SOP using different versions of operating systems and R are shown in **Table 1a.**

**Table 1a.**
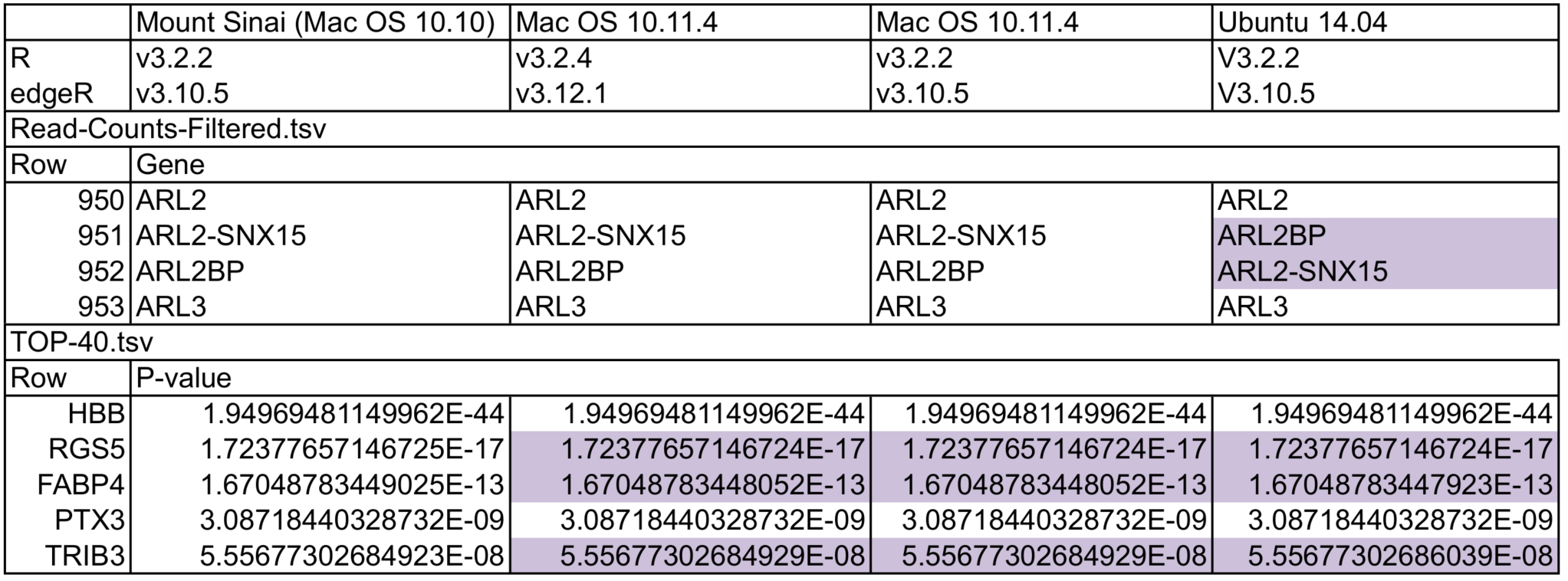
Reproducibility results comparing the expected results from Mount Sinai with results from two different operating systems with highlighted cells indicating a deviation. The differences in the output of 40 differentially expressed genes induced by a drug treatment (Trastuzumab) with the most significant p-values (“`Top–40.tsv`”) show how floating point calculations are not exactly the same. The output file from the alignment step (“`Read-Counts-Filtered.tsv`”) lists the counts of each transcript expressed across the samples. The ordering of transcripts in “`Read-Counts-Filtered.tsv`” is **not** guaranteed to be the same across different environments, creating downstream reproducibility issues where ordering is important.

BWA is designed to map sequence reads against a large reference genome and allows for mismatches and gaps. The input read sequences were generated by DToxS using the UMI protocol and were formatted as FASTQ files. BWA outputs the obtained alignment in the standard SAM (sequence alignment/map) format. A custom script from DToxS translates the SAM files into counts of transcripts, and the generated file “`Read-Counts-Filtered.tsv`" is one of the output files from the alignment step. This file lists the counts of each gene expressed across the samples. We experimented with Ubuntu, different versions of the operating system Mac OSX, R and the edgeR Bioconductor package (see **Table 1a**). We observe that the Ubuntu environment ordered the genes differently than Mac OSX. This difference in sorting is potentially problematic if the next stage processes gene counts in blocks, as splitting the genes at the points of discrepancy would lead to gene blocks that depend on the operating system. It is highly unlikely that academic researchers consider these types of discrepancy issues across operating systems when creating new bioinformatics tools.

The file “`Top-4 0.tsv`” shows the output from the differential expression analysis stage, where the top 40 differentially expressed genes induced by a drug treatment (Trastuzumab) in cardiac-derived cells with the most significant p-values across the samples are listed. We observe differences in floating point calculations of the p-values even with the same version of R and edgeR when the version of Mac OSX is changed (see **Table 1a**). While the differences in floating point values were not large enough to change the ordering of genes, these small differences may be amplified as more analysis modules are added to the workflow, each outputting slightly different floating points.

After observing the reproducibility from following each of the two steps from the DToxS SOP, we created two Docker containers, one for the alignment step, and a second for the differential expression step (see **Figure 1**). After creating the Docker containers and images, the two containerized steps of the DToxS workflow were run across different operating systems and computing environments so as to compare the outputs to the published results from DToxS. Specifically, we executed the containerized workflow on Mac OS, Microsoft Windows 10, Ubuntu, and the Google Cloud Container Engine. Table 1b shows that the containerized workflow yielded completely identical outputs across all environments that we tested.

**Table 1b.** Reproducibility results on different operating systems using Docker. Here, the results are exactly the same no matter what operating system is used to run the Docker containers.

|  | Docker on MacOS | Docker on Windows 10 | Docker on Ubuntu | Docker on Google Cloud |
| --- | --- | --- | --- | --- |
| Read-Counts-Filtered.tsv |  |  |  |  |
| Row | Gene |  |  |  |
| 950 | ARL2 | ARL2 | ARL2 | ARL2 |
| 951 | ARL2BP | ARL2BP | ARL2BP | ARL2BP |
| 952 | ARL2-SNX15 | ARL2-SNX15 | ARL2-SNX15 | ARL2-SNX15 |
| 953 | ARL3 | ARL3 | ARL3 | ARL3 |
| TOP-40.tsv |  |  |  |  |
| Gene | P-value |  |  |  |
| HBB | 1.94969481149962E-44 | 1.94969481149962E-44 | 1.94969481149962E-44 | 1.94969481149962E-44 |
| RGS5 | 1.72377657146724E-17 | 1.72377657146724E-17 | 1.72377657146724E-17 | 1.72377657146724E-17 |
| FABP4 | 1.67048783447923E-13 | 1.67048783447923E-13 | 1.67048783447923E-13 | 1.67048783447923E-13 |
| PTX3 | 3.08718440328732E-09 | 3.08718440328732E-09 | 3.08718440328732E-09 | 3.08718440328732E-09 |
| TRIB3 | 5.55677302686039E-08 | 5.55677302686039E-08 | 5.55677302686039E-08 | 5.55677302686039E-08 |

In summary, we show that the results of RNA-seq data analyses are dependent on the computing environment. These differences result in non-reproducible sorting of differentially expressed genes. Diagnostic tests often depend upon a profile of the top N differentially expressed genes, and even minor differences in sorting have the potential to significantly affect the final results. To address this issue, we show that software containers can be used to ensure the reproducibility of bioinformatics workflows by encapsulating the entire computing environment and software dependencies.

## Methods

### Generation of RNAseq data using the UMI protocol

The DToxS Center generates mRNA-sequencing datasets through the following three steps:

#### Cell Culture

Multiple samples of primary adult human cardiomyocytes (using a commercial cell line PromoCell) are subcultured and treated with different cancer drugs. After 48 hours, total RNA is extracted and transferred to a 96/384-well PCR plate for all cell samples. (SOP 1-4)

#### Library Preparation

The RNA sequence libraries are prepared according to the Single Cell RNA Barcoding and Sequencing method^18^ by converting Poly(A)+ mRNA sequences to cDNA sequences decorated with universal adapter, sample barcode and unique molecular identifier (UMI)^19^. (SOP 5)

#### cDNA Sequencing

The constructed cDNA libraries are sequenced by Illumina HiSeq 2500 platform and the output sequence datasets are de-multiplexed into a set of customized paired-end sequence data files in FASTQ format: the first read file includes a 10-bp UMI tag and a 6-bp sample barcode, and the second read file contains 46-bp cDNA sequence fragment. (SOP 6)

#### Data Analysis

The cDNA sequence fragments contained in the paired-end FASTQ data files are first aligned with the human genome reference library to quantify genome-wide gene expression levels for all cell samples, and then the gene expression levels between control condition and drug-treated condition are compared for each gene to identify the differentially expressed genes (DEGs) whose expression levels are changed significantly due to drug treatment. (SOP 7-8)

### Standard Operating Procedures (SOPs)^12^

1. DToxS SOP CE – 1.0: PromoCell Cardiomyocyte Subculture
2. DToxS SOP CE – 2.0: PromoCell Cardiomyocyte Plating for Drug Test
3. DToxS SOP CE – 4.0: Drug Treatment and Cell Lysis
4. DToxS SOP A – 1.0: Total RNA Isolation
5. DToxS SOP A – 6.0: High-throughput mRNA Seq Library Construction for 3' Digital Gene Expression (DGE)
6. DToxS SOP A – 7.0: Sequencing 3'-end Broad-Prepared mRNA Libraries
7. DToxS SOP CO – 3.1: Generation of Transcript Read Counts
8. DToxS SOP CO – 4.1: Identification of Differentially Expressed Genes

#### Software availability

We created Docker containers for each of the alignment stage and the differential expression stage in the DToX SOP. This was done by creating Dockerfiles that describe the dependencies and commands used to run each stage. The Dockerfiles were then used to build a Docker image, which is stored on the Docker Hub (https://hub.docker.com/u/biodepot/) and is publicly available for download. Both Docker images start with a base Ubuntu image and then install the necessary dependencies. The Dockerfiles are available from https://github.com/BioDepot.

## Acknowledgements

L.H.H. and K.Y.Y. are supported by NIH grant U54HL127624. Y.X. and E.S. are supported by NIH grant U54HG008098.

## Author contributions

T.M. implemented the Docker containers and conducted the reproducibility experiments. K.Y.Y. drafted the manuscript. K.Y.Y. and E.S. designed and coordinated the study. L.H.H. tested and refined the containers. Y.X. developed the computational analysis pipeline at DToxS. Y.X. and E.S. wrote the SOPs. All authors edited the manuscript.

